# Linear motor driven-rotary motion of a membrane-permeabilized ghost in *Mycoplasma mobile*

**DOI:** 10.1101/286278

**Authors:** Yoshiaki Kinosita, Makoto Miyata, Takayuki Nishizaka

**Affiliations:** Department of Physics, Gakushuin University, 1-5-1 Mejiro, Toshima-ku, Tokyo 171-8588, Japan; Graduate School of Science, Osaka City University, 3-3-138 Sugimoto, Sumiyoshi-ku, 8 Osaka 558-8585, Japan

## Abstract

*Mycoplasma mobile* exhibits a smooth gliding movement as does its membrane-permeabilized ghost model. This exceptionally prominent experimental system has allowed us to conclude that the energy source for *M. mobile* motility is adenosine triphosphate (ATP), and the gliding is largely comprised of repetitions of unitary steps of about 70 nm. In the present study, we show a new motility mode, in which the ghost model prepared with a high concentration of detergent exhibits directed rotational motions with a constant speed. With a rotational speed and viscous friction of a single ghost, the torque was estimated to be ∼30 pN nm at saturated [ATP]s. Although the origin of the rotation has not been conclusively settled, we found that rotary ghosts treated with sialyllactose, the binding target for leg proteins, were stopped. This result suggested that biomolecules embedded on the cell membrane nonspecifically attaches to the glass and works as a flexible pivot point, and the linear motion of the leg is a driving force for a rotary motion. This simple geometry exemplifies the new mechanism, by which the movement of a linear motor is efficiently converted to a constant rotation of the object on a micrometer scale.

---

*Mycoplasma mobile* (*M. mobile*) is a flask-shaped bacterium under optimal conditions that can smoothly glide on a solid surface in the direction of a protrusion at a speed of up to 4.5 μm s^-1^ [1] (Fig. 1A *top*). It lacks genes encoding conventional motor proteins such as myosin and kinesin, or bacterial flagella [2]. Three proteins essential for gliding are localized at a cell pole, and named as Gli123, Gli349 and Gli521 [3, 4]. These proteins are organized on the surface of machinery, and each of their numbers is estimated to be ∼ 450 (Fig. 1A *bottom*). Their molecular functions are as follows: Gli123 is a scaffold for other molecular machineries [5]; Gli349 binds to sialylated oligosaccharides (SOs) on a solid surface as a leg [6, 7] [8]; and Gli521 transmits the force into the leg as the crank [9, 10]. The internal structures, including the α-and β-subunit homologs of F-type ATPase, co-localized on the gliding machineries, suggesting that the internal structure might function as the motor for *Mycoplasma* gliding [11, 12]. Notably, the membrane-permeabilized ghost model revealed that the gliding machineries are driven by ATP hydrolysis; additionally, the smooth gliding movement was comprised of the repetition of 70-nm unitary steps [13, 14]. Although the actors in the motility have been gradually clarified, the mechanism by which the motor converts chemical energy into mechanical work remains an open question.

**Figure 1.**
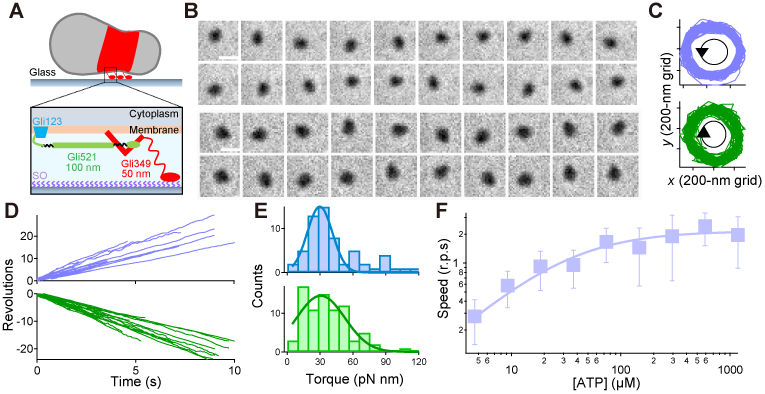
ATP-dependent rotation of tethered ghosts. (A) Schematics of a single cell (*top*) and a gliding machinery (*bottom*). (B) Upper and Bottom: Sequential images of a ghost in CCW and CW rotation at 100-ms intervals, respectively. Scale bar,1 μm. (C) The trajectories of the center of mass in CCW and CW rotation. Blue and green represent CCW and CW rotation, respectively, and their colors coincide with d and e. (D) Time course of revolution of the tethered ghost. (E) Histograms of a torque in both directions. Torque was calculated by a rotational speed and a viscous friction in each cell. Solid lines represent the Gaussian function, where 30.7 ± 21.9 and 29.8 ± 11.5 pN nm in CW and CCW rotations, respectively (n = 84 in CW rotation, n = 66 in CCW rotation). (F) The rotation rate of ghost at different [ATP]s (n = 480). Solid line showed the Michaelis-Menten kinetics: *V* = *V*_max_ [ATP]/(*K*_m_+[ATP]), where *V*_max_ and *K*_m_ were 2.2 Hz and 32 μM, respectively.

To explore the motor function in greater detail, we here constructed a motility assay that enabled detection of the rotary ghosts. Normally, the membrane-permeabilized ghost model is prepared with 0.09% Triton X-100, and the large number of ghosts showed a gliding motion after addition of ATP [13, 14]. In contrast, we found that a small percentage of ghosts prepared with a 0.013% concentration of detergent rotated at a fixed position like tethered-flagellated bacteria [15-17] (Supplementary Movie 1). In this condition, we also detected the gliding motion in ∼50% of the ghost population, and the gliding speed was similar to that of live cells at saturated [ATP]s (Fig. S2).

Rotational motions occurred in both directions, and the population of each cell was 56% in the CW and 44% in the CCW direction (n = 150; Fig. 1B). Note that the center position of the rotation did not move and the radius also remained constant, indicating that some flexible part was connected to a glass surface (Fig. 1C). Rotational rate was calculated from the slope of the revolution (Fig. 1D). With the rotational rate and viscous friction of ghost (see Method section), the frictional torque against the surrounding solution was estimated to be 30.7 ± 21.9 in the CW direction, and 29.8 ± 11.5 pN nm in the CCW direction at a saturated [ATP] (n = 84 in CW, n = 66 in CCW; Fig. 1E). We did not see a difference in the CW and CCW directions, and therefore analyzed CW and CCW rotation as the same (*P* = 0.1983>0.05 by *t*-test).

We next investigated the effect of ATP concentration on the rotational rate of ghosts. In the range of 4–1200 μM [ATP]s, the relationship between a rotational rate and [ATP] obeyed simple Michaelis-Menten kinetics, where *V*_max_ and *K*_m_ were 2.2 Hz and 32 μM, respectively (n = 480; Fig. 1F). This suggested that the motor had no cooperativity in the ATP binding event. Note that the discrete steps were detected under low [ATP]s, suggesting that ATP binding is the rate-limiting step for rotary movement in this condition (Fig. S3); in addition, the high-speed imaging enabled us to detect stepwise motions even at saturated [ATP]s. From this analysis, step size, stepping torque and rate limiting steps were estimated to be 34°, 85–120 pN nm, and 15.5 s^-1^, respectively (Supplementary Result and Discussion).

To investigate whether the leg protein contributed to the rotation, we treated ghosts with sialyllactose (SL), a binding target for legs [18, 19] (Fig. 2A). Notably, rotary ghosts did not detach from the glass surface but slowed down and/or stopped, while gliding ghosts detached from the surface (Supplementary Movie 2, Fig. 2B). Additionally, the ratio of rotational rate (*f*_after_/*f*_before_) decreased in proportion to [SL] (Fig. 2C). In this condition, the discrete stepwise movements were also detected even at saturated [ATP]s, which might correspond to the binding time of the leg to the SO (Fig. 2D). To determine the stepping angle quantitatively, we next fitted the data with a step-finding algorithm (see Methods section). From this analysis, the step size was estimated to be 33.1 ± 10.1°, assuming that the histogram comprises a single peak (Fig. 2E; n = 125). The dwell time between steps depended on [SL], where the average and SD were 0.37 ± 0.39 in 0.5 mM and 0.55 ± 0.51 s in 3 mM, respectively (n = 83 in 0.5 mM, n = 71 in 3 mM; Fig. 2F). This result suggested that the leg binds to free SL and consequently, could not produce the thrust for rotation.

**Figure 2.**
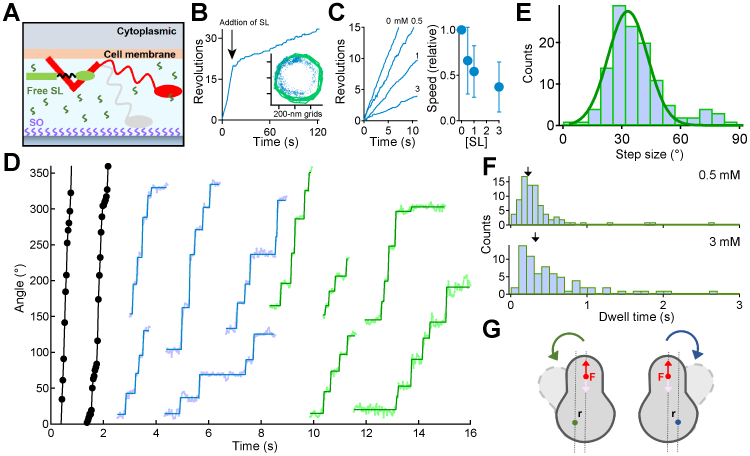
Linear movement generated the rotary motion. (A) Schematic illustration of the experiment. Gray and red legs are before and after addition of the sialyllactose (SL), respectively. (B) Typical example of the time course of revolution in presence of a free SL in solution. The arrow represents the time when SL were added. *Inset*; Blue dot and green line at 33-ms intervals represent a rotational trace before and after addition of the free SL in solution, respectively (C) Dependence of a free SL on a rotation rate at the saturated [ATP]. *Left*; Time course of revolution under the various [SL]s, 0.5-3 mM. *Right*; The relationship between the rotational speed and mM [SL] (n = 56). The relative speed *f* represents the *f*_after_/*f*_before_. (D) Typical examples of a stepwise rotation under the various [SL]s. Black, cyan, and green dots represent the raw data at 0, 0.5, 3 mM [SL], respectively. Rectangles in the rotation are lines fitted by the step-finding algorithm (see the methods section in detail). (E) Histogram of the step size calculated by a step-finding algorithm (n = 125). Solid line showed the Gaussian function, where the size was 33.1 ± 10.1 degree. (F) Histograms of dwell times at 0.5 and 3 mM [SL]. Arrows in each graph indicate the average value which are 0.37 ± 0.39 in 0.5 mM and 0.55 ± 0.51 sec in 3 mM, respectively (n = 83 in 0.5 mM, n = 71 in 3 mM). (G) Schematics of the pivoting model for rotation. The green and blue dots represent the tethered point to the surface. The red dot represents the position of active leg(s); additionally, the red and pink arrows correspond to the direction of thrust from a surface and the direction of leg’s force to a surface, respectively. The width of gray-dot lines (r) represents the distance between the power stroke of the active leg and tethered point. Depending on the tethered point, the rotational direction could be changed; e.g., the CW direction in the case of the blue tethering point.

We next explored the effect of the antibody on the rotation. We used monoclonal antibodies (MAb) MAb7 and MAbR19 against Gli349 and Gli521, respectively, which influence binding activity and gliding speed, respectively [20]. By treating ghosts with MAb7, the gliding ghosts dissociated from the glass surface, while the rotary ghosts stopped but did not detach from the glass. In MAbR19 experiments, both gliding and rotary ghosts stopped suddenly, but did not detach from the glass (Supplementary Movie 3). As stated previously, these results indicated that the crank Gli521 transmits the force, and the leg(s) Gli349 bind to and release from SO and produce the thrust [3, 4].

We considered possible mechanisms to explain the rotation. One scenario is the pivoting model: the leg produced the thrust for the rotation, while a flexible point such as the membrane is tethered to the surface (Fig. 2G). For a distance *r* between the power stroke of the active leg and tethered point to glass, the torque was estimated by the following equation: *T* = *r* × *F*, where *T* is the torque, and *F* is the thrust of leg [21]. With this model, bidirectional rotation could be produced depending on the geometry between the tethering point and the active leg. Given this model and assuming that a step length is 70 nm [13], the distance *r* was estimated to be 120 nm using the following equation: *L* = 2 *r* sin (*θ*/2), where *L* is a step length, and *θ* is a step angle (33°), which might correspond to the periodicity of gliding machineries.

Although the number of legs involved in rotation was not conclusively determined, the thrust could be estimated to be 0.7-1 pN from the above equation, assuming that the number of legs for rotation was driven by a single leg, which was similar to the values estimated by an optical tweezer [22]. Interestingly, the 120-nm length is consistent with the size of Gli521 [9]; therefore, another possible rotary model is that one part of the same gliding machinery, such as Gli521, was sticking to a glass surface while pulling another part of the complex such as Gli349. If the gliding machinery exhibits constant displacement and force, the various outputs will be detected depending on the geometry of the pivoting model, e.g., the detection of smaller output when r is small, and vice versa. Considering that the repetitive steps are clear, and the distribution of steps was narrow (Fig. S5), this model might also be possible.

In this study, we could not exclude the possibility that the rotation was driven by the internal structure of *M. mobile*, which is the homolog of F-Type ATPase and co-localized on the gliding machinery. This is because some ghosts had a rotation axis in the middle and rotated like a propeller, the centroid of which is at the middle of the ghost, suggesting that the real rotary motor was connected to the glass surface as shown in F_1_-ATPase and bacterial flagella [15-17, 23, 24] (Supplementary Movie 4 and Fig. S8A). Because the motor should rotate itself to maintain the cell body, actin or bead rotating, they could show fake rotation like the cowboy if the one end of tip or backward attached to a surface or γ-shaft. In *M. mobile*, however, the propeller rotation could also be explained by the pivoting mechanism.

To address whether *M. mobile* forms a rotary motor, 120°-steps should be detected in the propeller rotation. So far, we have detected step-like motions, though not 120°, at a few μM range of [ATP] (Fig. S8B). But we inferred that the next chemical cycle would start before the completion of mechanical work in this condition. Although we should improve the experimental condition to detect the rotation and steps at nM [ATP] like F_1_-ATPase [23], our novel assay would be helpful for clarifying the mystery of whether F-type ATPase is the motor for *Mycoplasma* gliding. Finally, as demonstrated in *Flavobacterium johnsoniae*, the molecular motor of which its rotary motion converts into linear motion might be common feature in gliding bacteria [25].

## Acknowledgements

We thank Richard Berry for discussions that were critical in preparing the manuscript and Mitsuhiro Sugawa for developing the step-finding algorithm.

## Author Contributions

Y.K. and M.M. designed research; Y.K. performed research; Y.K. and T.N. constructed the optical setup and microscope; Y.K., M.M., and T.N. wrote the paper.

