## Supplementary Materials for "Linear motor driven-rotary motion of a membrane-permeabilized ghost in *Mycoplasma mobile*"

This file includes:

Supplementary Methods

Supplementary Results and Discussion

Supplementary Figures 1-8

Captions for Supplementary Movies

### Supplementary Methods

#### Strains and cultivation

*M. mobile* strain (ATCC 43663) was grown in Aluotto medium [2.1% (wt/vol) heart infusion broth, 0.56% yeast extract, 10% (vol/vol) horse serum, 0.025% thallium acetate, and 0.005% ampicillin] at 25 °C until the absorbance at 600 nm reached 0.06–0.10 [1].

#### Tethered ghost assay

Preparation of ghosts was previously described except that the all buffer contains 0.1 % methylcellulose (M0512; Sigma Aldrich) to prevent ghosts from dissociating from a glass surface [2, 3]. The final buffer was comprised of 10 mM Tris-HCl at pH 7.5, 50 mM NaCl, 1 mM DTT, 1 mM EGTA, 2 mM $MgCl_2$, 0.5 mg/ml Bovine serum albumin, $Mg^{2+}$-ATP with an ATP-regenerating system (0.2 mg $ml^{-1}$ creatine kinase and 0.8 mg $ml^{-1}$ creatine phosphate) and without methylcellulose. To avoid the depletion of ATP, each experiment was done within 30 min. All experiments were done at RT.

To check the effect of the free sialyllactose (A0828; Sigma Aldrich) and antibodies on the rotation, the 20-ml volume of final buffer containing these reagents was induced into the chamber. The final concentration of antibodies was ~5 μg $ml^{-1}$.

#### Microscope

Tethered ghosts were observed under a phase-contrast microscope (IX71; Olympus) equipped with a 100× objective (UPLSAPO 100 with Ph and 1.4 N.A.; Olympus), a CCD camera (VCC-H1600 or LRH1540N; Digimo), a highly-stable customized stage (Chukousha), and an optical table (RS-2000; Newport) [4]. Ghost position was determined by a centroid fitting or ellipsoid fitting with 8-nm accuracy [2, 4, 5] (Fig. S1).

**Data analysis**

Sequential images were captured as 8-bit images with CCD camera at 4- or 33-ms temporal resolution and converted into a sequential TIF file without any compression.

The rotational torque against around a solution is estimated as follow equation: $T = 2\pi f\xi$, where $T$ is the rotational torque, $f$ is a rotational speed, and $\xi$ is a viscous friction. The viscous friction is, for the case of a spherical-shaped ghost, given by $8\pi\eta a^3+6\pi\eta ar^2$, where $\eta$ is the viscosity, $a$ is the radius of the ghost, and $r$ is the radius of rotation. We frequently observed the ellipsoidal-shaped ghost like two spherical-ghost combined. In this case, the viscous friction is given by $2\times 8\pi\eta a^3+6\pi\eta ar_1{}^2+6\pi\eta ar_2{}^2$, where $r_1$ and $r_2$ are the radii of the revolution of the inner and outer ghosts, respectively [6]. The radius of a ghost was measured using ImageJ 1.48v software (http://rsb.info.nih.gov/ij/). The average viscous friction was estimated to be 2.4 pN nm s.

To determine the stepping angle quantitatively, we used two processes as follows. To roughly extract more four steps with a regular size of steps, pairwise distance function analysis (PDF analysis) was first applied [2]. We analyzed 129 ghosts at the various [ATP]s, and 267 runs in 44 ghosts were shown more than four steps. Typical examples of PDF analysis were shown in Fig. S4. Next, to extract the instantaneous stepping motion, we applied the step-finding algorithm to 44 ghosts [2, 7]. Because the instantaneous stepping motion was only fitted with a step-finding algorithm, we inferred that the multiple legs-driven motion could be excluded in this analysis. A step-finding algorithm was applied to 267 runs in 44 ghosts, and 178-time courses in 40 ghosts were successfully extracted (Fig. S4).

For estimation of stepping rate between steps, we select the 30°-40° steps and averaged the consecutive steps for 0.2 sec [8]. Time zero for each step record was assigned closest to 15°. To eliminate the contribution of short pauses, we applied the linear fitting to the averaged step between 4° and 30°, where the slope corresponded to the stepping rate.

### Supplementary Results and Discussion

#### 1. High-speed imaging of stepping rotations under various [ATP]

We successfully detected the stepwise rotation under low [ATP]s and in the presence of sialyllactose. To gain mechanical insight into the motility mechanism under various [ATP]s, we next performed high-speed imaging of rotations with a time resolution of 4 ms. Remarkably, we also detected the intermittent pauses at saturated [ATP]s (Fig. S4). The 178-time courses in 40 ghosts showed the instantaneous stepping motion, and could be successfully fitted by a step-finding algorithm (outputs are superimposed as a line in Fig. S4). We could not see any differences of step angles in the range of 11–347 μM [ATP] (Fig. S5). The histogram of step angles under various [ATP] comprised of a single peak, where the peak and SD were 34.1±19.6°, respectively (Fig. S5; n = 842).

#### 2. Stepping rotations

With the stepping rate and viscous friction of a single ghost, the stepping torque *T* under various [ATP]s could be estimated as the following equation: *T* = the stepping rate × the viscous friction of ghost (2.4 pN nm s). To check whether the stepping torque was dependent on [ATP], we then investigated the stepping rate. We extracted consecutive steps of 30-40º from the rotational record at each [ATP], and each step was averaged and superimposed as the thick cyan line in Fig. S6. To eliminate the contribution of short pauses, we calculated the slope between 4–30º with a linear fitting. From this analysis, the stepping rate was estimated to be 35-50 rad $s^{-1}$ under various [ATP]s (Fig. S6). Based on the fact that the ghost's shape was the same under various [ATP]s, this result indicated that ghosts produced constant torque irrespective of [ATP]. From this analysis, the stepping torque was estimated to be 85–120 pN nm, which was a few times larger than

$F_1$-ATPase [9], but 10 times smaller than the bacterial flagellar motor [10].

#### 3. Dwell-time analysis

To realize the molecular mechanism in terms of chemo-mechanical coupling, we next extracted the dwell time with the step-finding algorithm. At [ATP] < $K_m$ = 32 μM, the distribution of dwell time showed double exponential decay, indicating that the stepwise motion was comprised of two chemical reactions. To determine the rate constant of individual reactions, histograms were fitted with a double exponential: const·(exp(-$k_1$×t)-exp(-$k_2$×t)). In the range of 11-44 μM ATP, $k_1$ was increasing in linear proportion to external [ATP]s, indicating that $k_1$ was the rate of ATP binding (pink circles in Fig. S7). Even at [ATP]> $K_m$, histograms showed a double exponential decay, suggesting that two reactions of ATP-independent manners could exist, such as the Pi release and binding time of Gli349 to the SO. Histograms were fitted with a double exponential at [ATP] >$K_m$, where two rate constants were around 30 $s^{-1}$, which was similar to the value of the ATP-independent rate at low [ATP] (green circles in Fig. S7). To describe the chemical reaction at all [ATP]s, global fitting was applied to all histograms: const·(exp(-$k_{on}$×[ATP]×t)-exp(-$k_1$×t)) at 11–44 μM ATP and const·(exp(-$k_1$×t)-exp(-$k_2$×t)) at 87–347 μM ATP, where $k_{on}$ for ATP-dependent reactions, and $k_1$ and $k_2$ for ATP-independent reactions were calculated to be $6.7\times10^5$ $M^{-1}\cdot s^{-1}$, 29.6 and 31.9 $s^{-1}$, respectively. The rate constant of ATP-independent reactions was similar to a previous report, but the ATP-binding rate was 20 times smaller [2]. This discrepancy might be due to the reduction of active legs for the movement treated with a high concentration of detergent. This interpretation coincides with reports on other molecular motors where dwell time was proportional to the number of active motors [11].

#### 4. Difference between averaged torque and stepping torque

By applying high-speed video microscopy for observation, we also detected a 34°-stepwise movement and calculated the stepping rate as 35-50 rad $s^{-1}$, which corresponded to 6–8 Hz. In the dwell-time analysis, the ATP-independent two rate constants were estimated to be ~30 and 32 $s^{-1}$, respectively, indicating that the rate-limiting steps would be 15.5 $s^{-1}$. The average rotational rate was thus calculated with the following equation: $V$ = Step size × rate = 34°× 15.5 $s^{-1}$ = 527 ° $s^{-1}$, corresponding to 1.5 Hz, which roughly coincided with the maximum speed calculated by Michaelis-Menten kinetics (Fig. 1F). We thus concluded that the above discrepancy was due to including the dwell time in the analysis in the video-rate observation.

#### 5. Energy-conversion efficiency

Although we couldn't conclude that the number of gliding units are involved in the rotation, we inferred the possible motility mechanism of *M. mobile*. From the result that the ATP-dependent rate roughly showed a linear-dependent manner, each step might be comprised of a single turnover of ATP (Fig. S7). Assuming that one ATP molecule is consumed per step, the mechanical work can be estimated to be 50-70 pN nm with the following equation: $W$ = step size (34°) × stepping torque (85-120 pN nm). With this 1:1 scenario, the energy-conversion efficiency was roughly 50-70%, which supplies chemical energy of 100 pN nm when hydrolyzed.

#### Supplementary figures

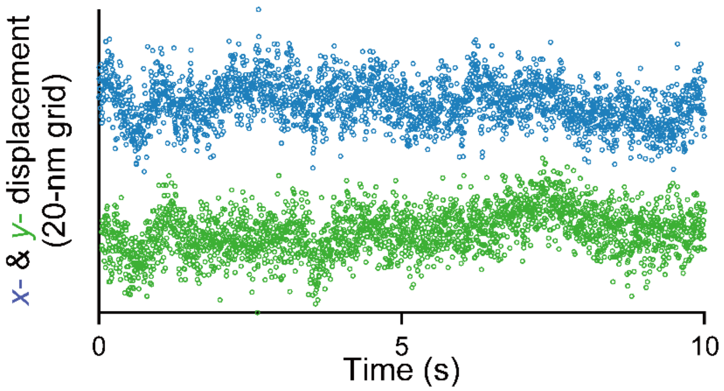


**Supplementary Figure 1 Stability and spatial resolution of our setup**

Immobilized ghost on a glass surface was captured with a temporal resolution of 4 ms under a phase-contrast microscope. The position was determined by a centroid fitting. The SDs of $x$ and $y$ were 7.7 and 7.5 nm, respectively, which were corresponded to a spatial resolution of our setup (n = 4).

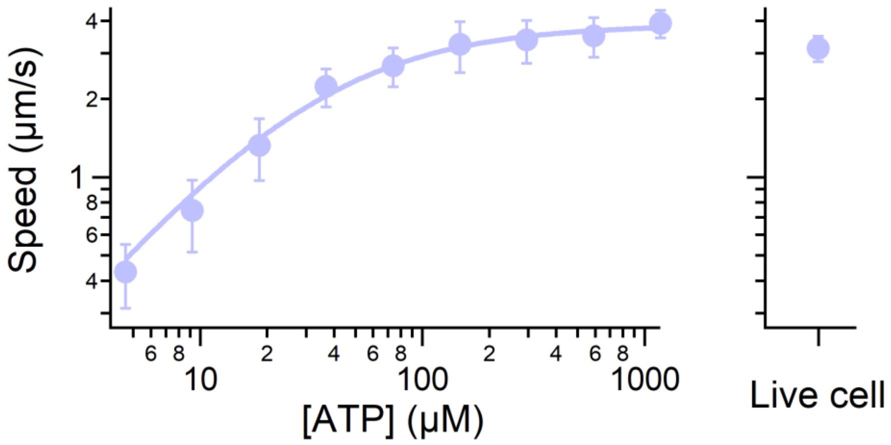


**Supplementary Figure 2 Gliding speed of ghosts at different [ATP]s**

*Left*; Data points represent average gliding speeds of ghosts under various [ATP]s (n = 172). Solid line shows $V=V_{max}[ATP]/([ATP]+K_m)$, where $V_{max}$ = 3.9 μm $s^{-1}$ and $K_m$ = 32 μM, respectively. *Right*; The speed of live cells (3.1±0.4 μm $s^{-1}$, n = 20).

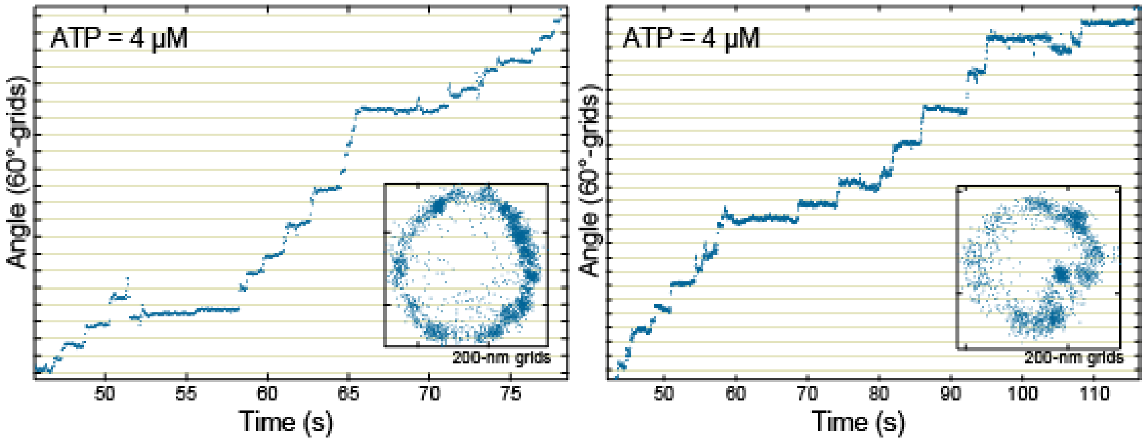


**Supplementary Figure 3 Stepwise rotation at low [ATP]**

*Inset*: *x*-*y* trace.

.

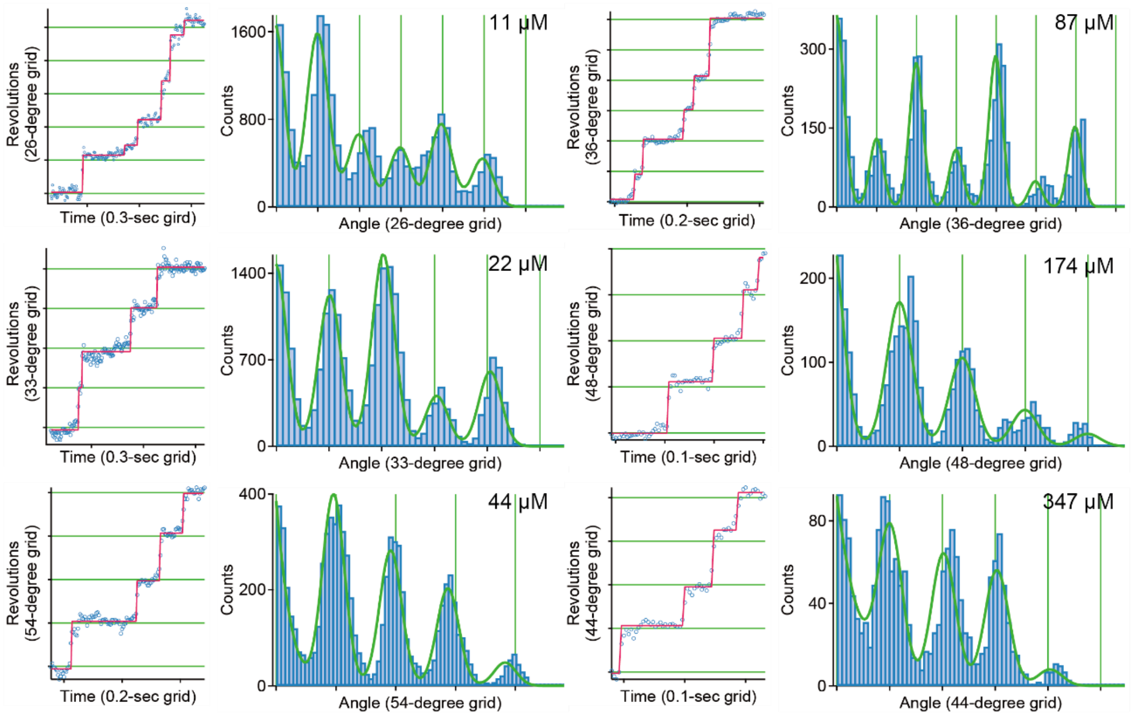


**Supplementary Figure 4 Time courses of stepwise rotations under various [ATP]s**

*Left*: The red line represents the fitting of the raw data by a step-finding algorithm. *Right*: Histograms of the pairwise distance function of the raw data. Green solid lines represent the sum of Gaussians.

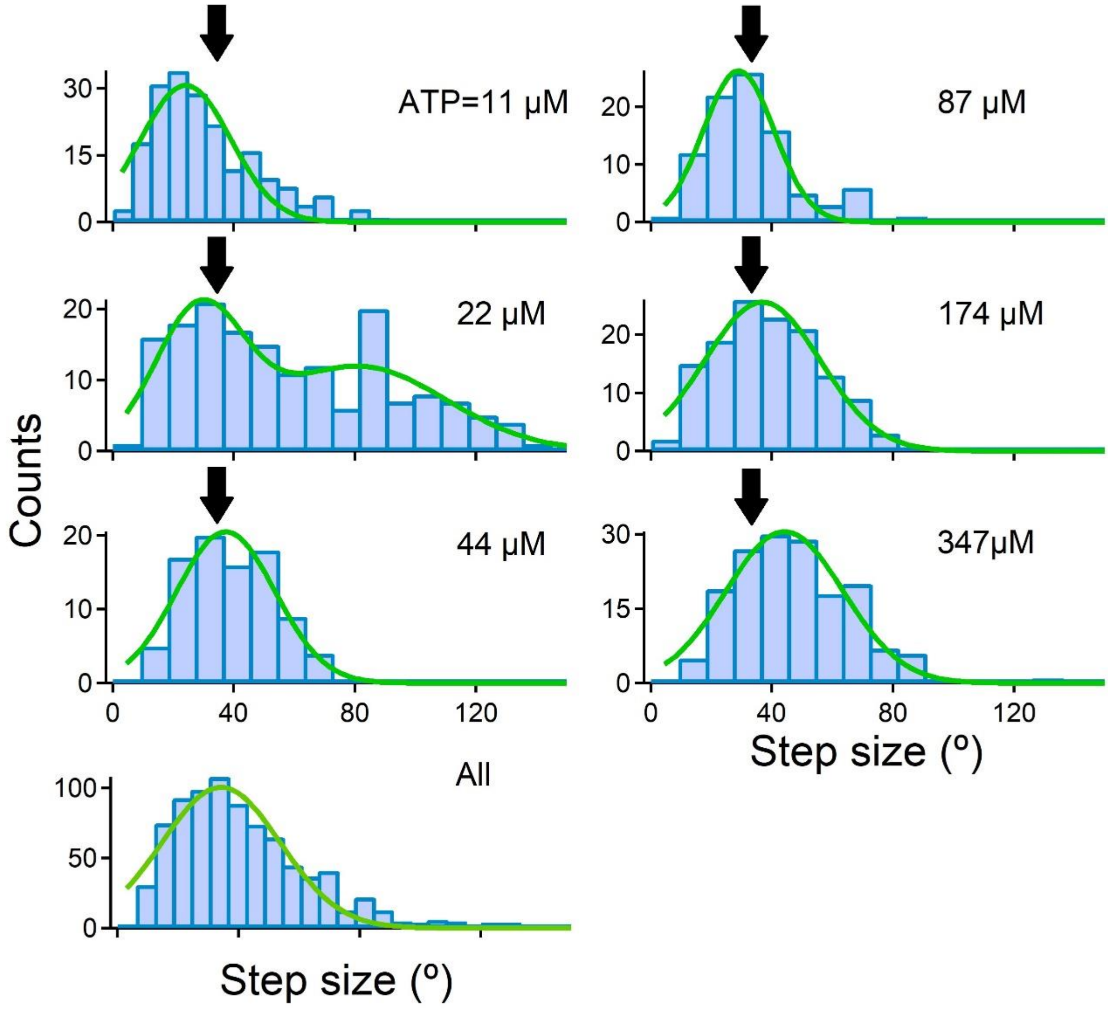


**Supplementary Figure 5 Histograms of the step angle extracted by a step-finding algorithm under various [ATP]s**

Solid lines represent the single or the sum of Gaussian(s). Allows indicate the average value of all data with a size of 34° (n = 841). The number of steps is 198, 169, 89, 92, 131, 162 at 11, 22, 44, 87, 174, 347 μM, respectively.

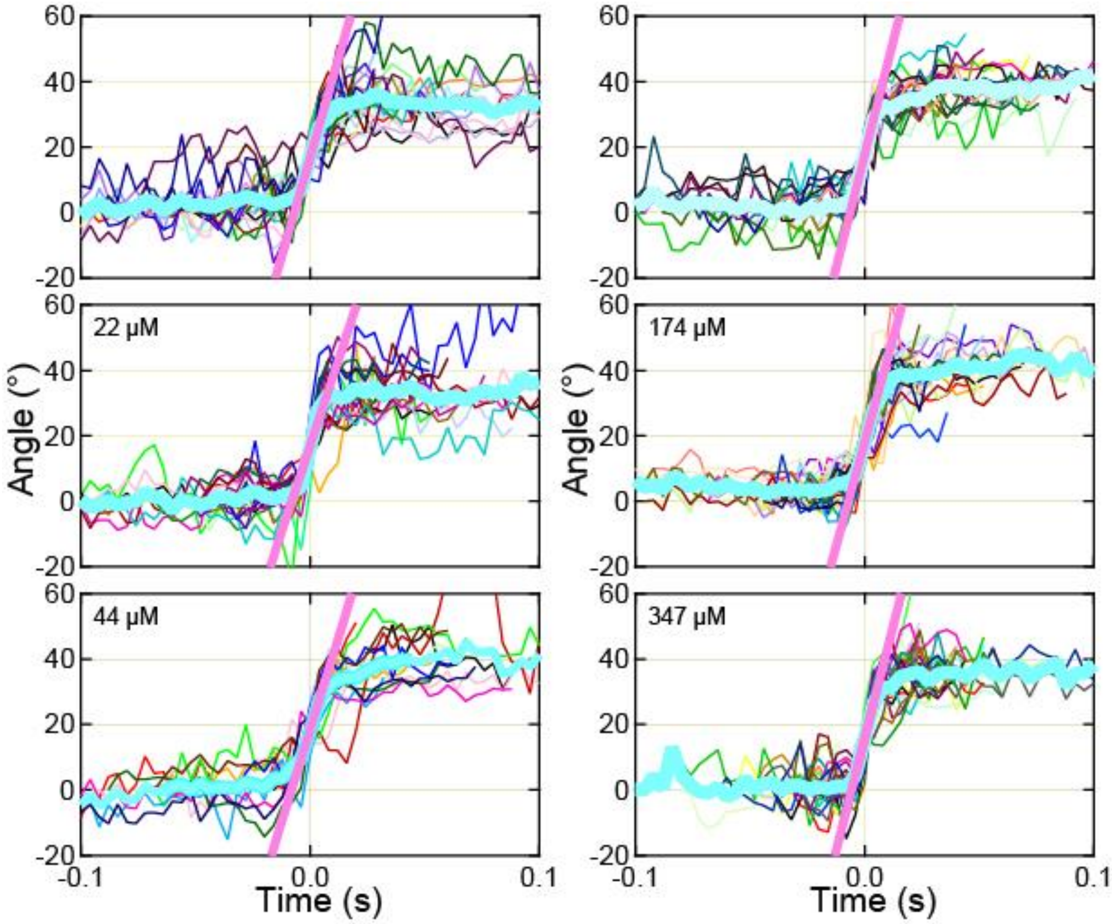


**Supplementary Figure 6 Stepping torque**

Stepping rate for torque estimation. Thin colored curve represents the raw data with the size of 30-40°steps; thick blue curves are their average (n = 26, 21, 18, 25, 26, 30 consecutive steps at 11, 22, 44, 87, 174, 347 μM [ATP]). Time zero for each step record was assigned closest to 15°. Pink thick lines represent the linear fit to each averaged trace between 4° and 30 °, providing the torque. The slopes at 11, 22, 44, 87, 174, 347 μM [ATP] were 42.64, 36.92, 39.63, 48.55, 44.70, 48.2 rad $s^{-1}$, which were corresponded to the stepping rate. With the stepping rate and the viscous friction of the single ghost (2.4 pN nm s), the stepping torque were estimated to be 85-120 pN nm.

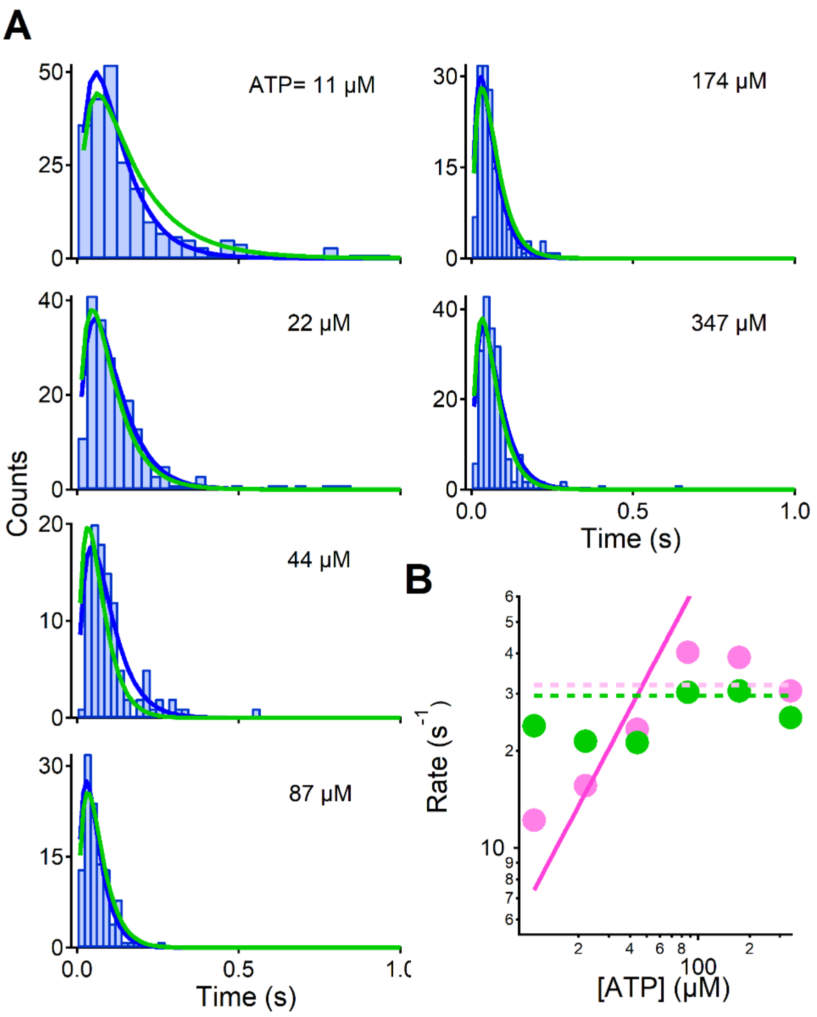


**Supplementary Figure 7 Dwell time analysis**

(A) Histograms of dwell times at various [ATP]s. Total counts are 233, 197, 105, 111, 156 and 193 steps in the range of 11- 347 μM [ATP]s. Blue lines are double exponential fits: constant·(exp( $-k_1\times$t)-exp($-k_2\times$t)), where $k_1$ and $k_2$ represent green and pink filled circle in B. Note that $k_1$and $k_2$ were changed above 44 μM ATP. Green line are global fitting: const.×(exp(-($k_{on}\times$[ATP]$\times t$)-(exp($-k_1\times t$) for 11-44 μM ATP and constant·(exp($-k_1\times$t)-exp($-k_2\times$t)) for 87-347μM ATP , where $k_{on}$ was the rate constant for ATP binding ($6.7\times10^5$ $M^{-1}$ $s^{-1}$) and two ATP-independent reaction with $k_1$ and $k_2$ were 29.6 and 31.9 $s^{-1}$ . (B) The relationship between rate constants and [ATP]. Green and pink dots represent the rate constants estimated by the exponential fit to individual data. Pink and green dot lines represent the rate constants obtained by the global fitting. Pink-solid line represents an ATP binding constant acquired with the analysis of a global fit.

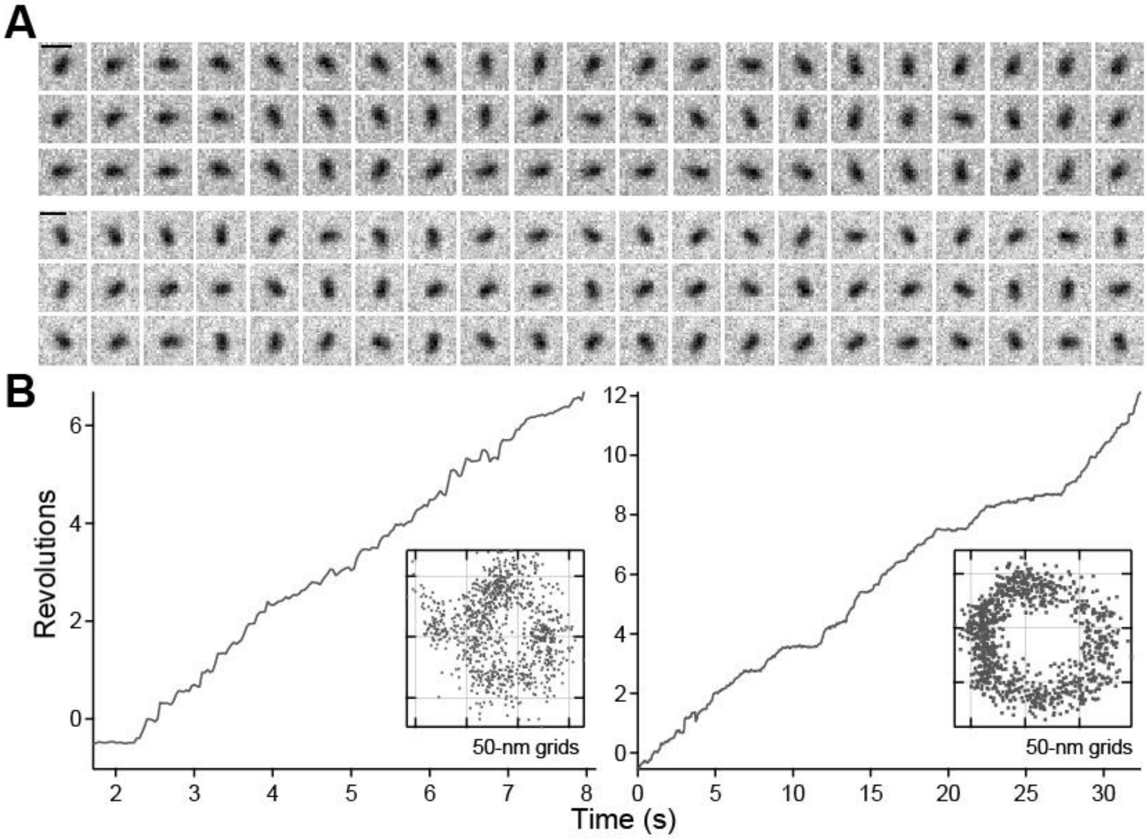


**Supplementary Figure 8 Propeller rotation**

(A) Sequential phase-contrast images of a propeller rotation at 100-ms intervals under 8 μM (*upper*) and 32 μM [ATP] (*lower*). The ghost had the rotation axis at the middle and rotated like a propeller Scale bar, 1 μm. (B) The time course of revolutions under 8 μM (*left*) and 32 μM [ATP] (*right*). *Inset*: *x*-*y* trace.

### Captions for Supplementary Videos

**Movie S1**

Rotational motion of ghost at a saturated [ATP] observed by a phase-contrast microscopy with a temporal resolution of 33 ms. Scale bar, 10 μm.

**Movie S2**

Inhibition of a rotational motion by sialyllactose. Sialyllactose was added into the buffer at about 10 sec. Scale bar, 10 μm.

**Movie S3**

Inhibition of a rotational motion by 0.5 mg $ml^{-1}$ antibody MabR19 which influenced on the crank protein, Gli521. Scale bar, 20 μm.

**Movie S4**

A propeller rotation at 8 μM [ATP]. The ghost had the rotation axis at the middle and rotated like a propeller Scale bar, 1 μm.
